# High-amplitude network co-fluctuations linked to variation in hormone concentrations over menstrual cycle

**DOI:** 10.1101/2021.07.29.453892

**Authors:** Sarah Greenwell, Joshua Faskowitz, Laura Pritschet, Tyler Santander, Emily G. Jacobs, Richard F. Betzel

## Abstract

Many studies have shown that the human endocrine system modulates brain function, reporting associations between fluctuations in hormone concentrations and both brain activity and connectivity. However, how hormonal fluctuations impact fast changes in brain network structure over short timescales remains unknown. Here, we leverage “edge time series” analysis to investigate the relationship between high-amplitude network states and quotidian variation in sex steroid and gonadotropic hormones in a single individual sampled over the course of two endocrine states, across a natural menstrual cycle and under a hormonal regimen. We find that the frequency of high-amplitude network states are associated with follicle-stimulating and luteinizing hormone, but not the sex hormones estradiol and progesterone. Nevertheless, we show that scan-to-scan variation in the co-fluctuation patterns expressed during network states are robustly linked with the concentration of all four hormones, positing a network-level target of hormonal control. We conclude by speculating on the role of hormones in shaping ongoing brain dynamics.

## INTRODUCTION

The human brain is a complex network composed of structurally connected neural elements that help shape brain activity and give rise to widespread patterns of functional coupling [1, 2]. The organization and topology of these structural and functional networks can be interrogated using tools from network science [3], revealing organizing principles including short processing paths [4], hubs and rich clubs [5, 6], modular structure [7], and cost-efficient spatial embedding [8].

Recent work has shown that whole-brain patterns of functional coupling between brain regions vary over short timescales [9–11]. To reconstruct changes in network structure over time, most studies use sliding-window analyses [12, 13]. In this approach, a functional network is estimated using only those samples that fall within a window of some fixed duration. The window is then advanced a certain number of frames, resulting in a time series of functional networks. Although applied widely, this approach has a number of drawbacks. Namely, it forces the user to specify parameters for window length and amount of overlap between successive windows. The windowing procedure, itself, also makes it impossible to precisely localize a network state to a specific moment in time and resolve changes in network structure over short timescales.

Recently, we proposed “edge time series” (ETS) as a method for decomposing functional networks into time-varying components [14–16]. This approach helps address some of the limitations of sliding-window analyses, in that it is parameter-free and can resolve changes in network structure at a framewise timescale. In previous studies, we used this method to show that fast network dynamics are not smooth, but rather are bursty, identifying long periods of quiescence punctuated by brief, network-wide, high-amplitude “events” [14, 17, 18]. These events are of particular interest, as time-averaged functional networks can be accurately reconstructed from a very small number of events. We also found that the patterns of co-fluctuation expressed during events contain disproportionate amounts of information about an individual and that events can improve brain-behavior correlations. Despite their apparent utility, the application of edge time series for linking brain dynamics to cognitive, clinical, or physiological phenomena has been limited [19, 20].

Edge time series are well-situated for investigating relationships between brain connectivity and physiological variables that also fluctuate over short timescales. A good example is the human menstrual cycle, which is typified by variations in the sex steroid hormones estradiol and progesterone and gonadotropins follicle-stimulating hormone (FSH) and luteinizing hormone (LH). Briefly, estradiol increases during the first phase of the cycle known as the follicular phase, encouraging growth of the uterine lining. Immediately prior to ovulation, FSH encourages an immature follicle to complete its development into an egg for release from the ovaries. LH promotes the release of the egg into the fallopian tubes, followed by an increase in progesterone during the so-called luteal phase, during which the uterine lining thickens, creating a favorable environment for the egg to be preserved. These hormones exhibit neuromodulatory effects, and their impact on brain activity has been well-documented. The impact of sex hormones on brain activity has been well-documented [21, 22]. However, most previous work has involved cross-sectional study designs with incomplete sampling of participants’ cycles and comprised of heterogeneous cohorts of individuals.

Recently, a series of “dense sampling” studies have characterized quotidian variation in these hormones and their relationships with functional connectivity and network community structure [23–26]. The design of these studies parallels that of other recent dense sampling studies [27–29], acquiring data from, in this case, a single individual over the course of two complete menstrual cycles, one in which the participant was naturally cycling and another while on an oral hormonal contraceptive regimen (referred to as Studies 1 and 2) [30]. Previous studies using these datasets have identified spatially-diffuse increases in functional connectivity and modular flexibility coincident with peaks in estradiol; progesterone, by contrast, was predominantly associated with reductions in connectivity [24, 26]. However, the dynamic underpinnings of these associations have not been fully explored.

Here, we investigate these same data, following the analysis pipeline from [18], in which we use edge time series to detect events from functional magnetic resonance imaging (fMRI) data and cluster whole-brain co-fluctuation patterns into a low-dimensional set of states. In agreement with previous studies, we find that events can be sub-divided into two distinct clusters. In particular, we show that the frequency with which the first cluster appears in a given scan is strongly correlated with quotidian variation in both FSH and LH. Furthermore we show that day-to-day variation in the edge- and system-level configuration of the first cluster is broadly associated with both FSH and LH, as well as progesterone and estradiol. Collectively, our results establish a new link between high-amplitude, network-level events and the human endocrine system, opening up new avenues for exploring the dynamic interplay between brain-hormone relationships.

## RESULTS

### Clustering high-amplitude co-fluctuations reveal distinct patterns of connectivity

Recent methodological advances have made it possible to track rapid fluctuations in functional network architecture, revealing the presence of “events” – short-lived and high-amplitude patterns of network-wide fluctuations [14–16]. The results of previous studies suggested that events are low-dimensional, such that a small repertoire of patterns are reiterated [17, 18]. Here, we test whether this was also the case in an independently-acquired dataset of a single individual.

To detect and assess whether there were repeated patterns of high-amplitude events, we used simple statistical procedure to identify sequences of temporally contiguous frames whose root summed square (RSS) exceeded that of a null model in which regions’ activity time courses were randomized (circular shifts). We retained only those sequences that were motion free, i.e. did not include a high-motion frame and were at least two frames away from any high-motion frame. For each motion-free sequence, we extracted the co-fluctuations expressed during the frame with the highest RSS value as a representative peak. In total, we detected 899 motion-free events (14.98 ± 5.27 events per scan session).

To detect putative “states,” “clusters,” or “communities” of events (note that here we use these terms interchangeably), we vectorized event co-fluctuation patterns and, for all pairs of events, computed their spatial similarity (Pearson correlation), resulting in a 899 × 899 matrix. We then used a variant of modularity maximization to assign each co-fluctuation pattern to a single cluster (see **Materials and Methods** for details). As in previous studies, we found evidence for two large consensus communities that appeared in most scan sessions and collectively accounted for ≈70.2% of all detected states (Fig. 2a). These clusters were highly reproducible across repeated runs of the modularity optimization algorithm (Fig. 2b) and divided event patterns into cohesive clusters (Fig. 2c). We note that these clusters were also robust to variation in processing pipelines, after splitting the data by experiment (see Fig. S3), and after systematically excluding data from individual scans in calculating the correlation (Fig. S5).

**FIG. 1.**
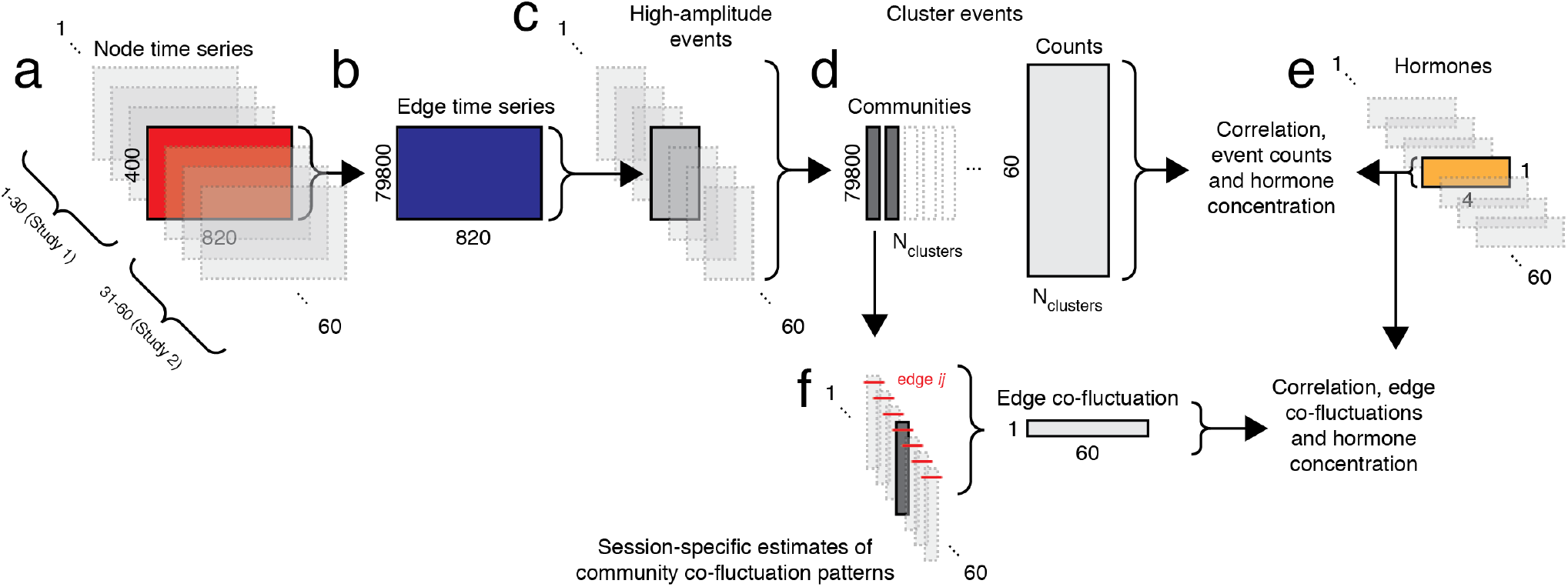
Analysis pipeline. (*a*) After preprocessing, we obtained parcellated regional time series from 60 scans (spanning two experiments). (*b*) For a given scan, we transformed node time series into edge time series following [14]. (*c*) Next, we detected high-amplitude events in each scan and for each event extracted its representative pattern (the frame with the greatest amplitude). In general, we obtained a different number of events per scan. (*d*) We aggregated event patterns from all scans and collectively clustered them using modularity maximization. This procedure resulted in multiple community centroids (we analyze the two largest) and a count of how many times a given community appeared on a given scan session. (*e*) In parallel, we analyzed hormone data that were collected concurrent with each scan session. Our principal aim was to link features of communities (brain states) with hormone data. (*f*) In addition, we reconstructed estimates of communities for each of the 60 scan sessions and, for each edge, computed the correlation of its co-fluctuation across sessions with hormone concentrations.

**FIG. 2.**
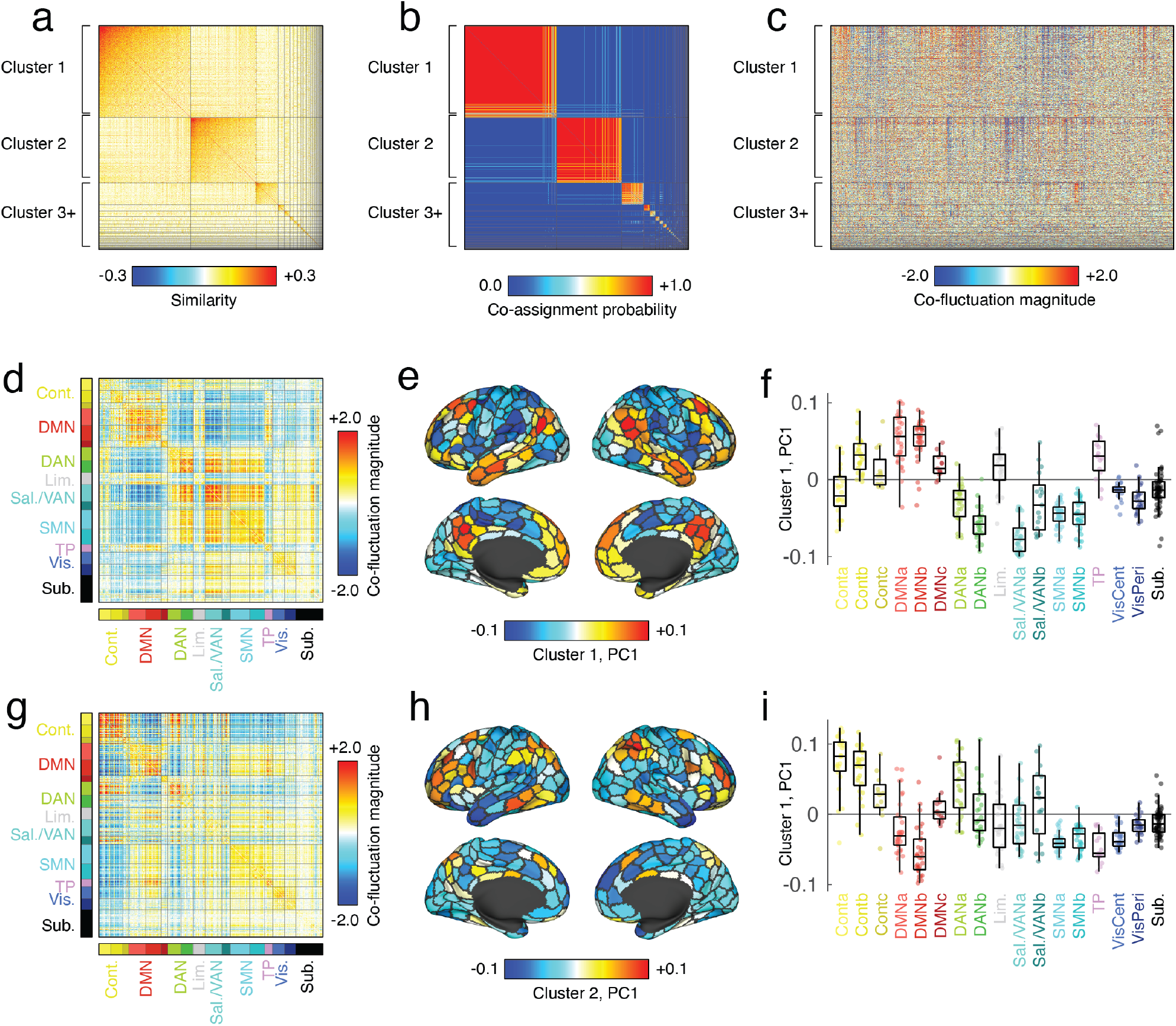
Modularity maximization and network states. We used an event detection algorithm to identify instances of “high-amplitude” co-fluctuations, extracting 899 brain-wide patterns. We calculated their spatial similarity and clustered them using modularity maximization. (*a*) Similarity matrix, ordered by communities (*b*) Community co-assignment matrix. (*c*) Vectorized co-fluctuation patterns ordered by communities. (*d*) Mean co-fluctuation matrix for cluster 1, ordered by canonical brain systems. (*e*) First principal component of the co-fluctuation matrix. (*f*) Elements of first principal component grouped by brain system. Panels *g*-*i* are analogous to *d*-*f* but for cluster 2.

Interestingly, the spatial patterns of these clusters closely recapitulate those reported in previous studies [14, 18]. Community 1, for instance, was characterized by opposed fluctuations between regions in the default mode network with those in dorsal and salience/ventral attention networks (Fig. 2d). To better understand whether these co-fluctuations were underpinned by a specific mode or pattern of activity, we performed a singular value decomposition of the mean co-fluctuation matrix, revealing, as expected, a node-level pattern characterized by strong fluctuations in default mode regions and opposed fluctuations in attentional and, to some extent, sensorimotor systems (Fig. 2e,f). Similarly, the spatial pattern of community 2 was characterized by opposed co-fluctuations of control and dorsal attention regions with the default mode network (Fig. 2g-i).

The remaining communities collectively accounted for < 30% of all events, with the next most frequent community accounting for 9.8% of all events (but appearing in ≈ 71.7% of scan sessions). For this reason, we focus on the first two communities for all subsequent analyses. In the supplementary material we describe the remaining communities in greater detail (Fig. S1).

**FIG. 3.**
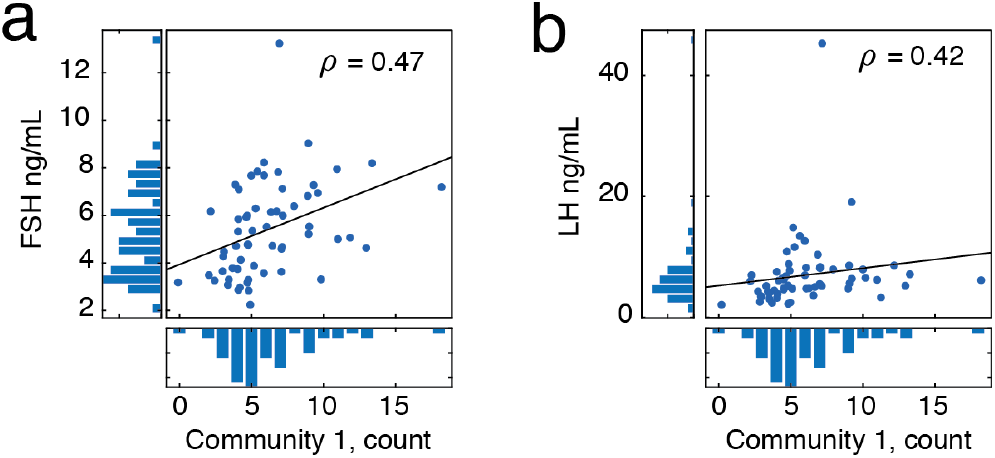
Correlations between state frequency and quotidian variation in gonadotropin concentration. (*a*) Scatterplot showing concentration of follicle-stimulating hormone across scan sessions *versus* the frequency with which cluster 1 appeared in a given scan. (*b*) Scatterplot showing concentration of luteinizing hormone across scan sessions *versus* the frequency with which cluster 1 appeared in a given scan.

### State frequency is associated with LH and FSH concentrations

Cluster analysis of co-fluctuation time series revealed the presence of repeating patterns or states. However, the biological relevance of these states remains unclear. In this section, we show that the frequency with which these states appear across scan sessions is robustly related to endogenous variation in the gonadotropins, luteinizing and follicle-stimulating hormone, but not statistically associated with sex hormones estradiol and progesterone.

To link high-amplitude co-fluctuations with quotidian variation in hormone concentration (see Fig. S2), we calculated for each scan session the number of times that each community (1 and 2) appeared. We found that, on average, communities 1 and 2 appeared 6.15 ± 3.17 and 4.37 ± 2.57 times (≈ 41% and ≈ 29% of all events; maximum values of 18 and 11), respectively. Next, we computed the Spearman rank correlation of these frequencies with hormone concentrations. Note that the rank correlation reduces statistical biases originating from spikes in both luteinizing and follicle-stimulating hormone around ovulation. For all reported correlations we corrected for multiple comparisons by fixing the false discovery rate at *q* = 0.05, resulting in an adjusted critical value of *p*_*adj*_ = 0.0067. We found statistically significant correlations between the frequency of community 1 with both gonadotropic hormones (*ρ*_1,*FSH*_ = 0.47 and *ρ*_1,*LH*_ = 0.42; *p* = 0.0002 and *p* = 0.0007). Interestingly, the correlations between community 1 and progesterone (*ρ*_1,*P*_ = − 0.33; *p* = 0.007 and community 2 with luteinizing hormone (*ρ*_2,*LH*_ = 0.26; *p* = 0.02) were both statistically significant at uncorrected levels, but failed to pass after multiple comparison corrections (see Fig. S6). Note that these results hold after splitting the 60 scans by experiment (Fig. S4) and with an alternative processing pipeline that does not include global signal regression (Fig. S7).

Collectively, these results suggest that endogenous and exogenously-induced changes in gonadotropic hormone concentrations are related to the expression of distinct brain network states. These findings posit a hormonal basis for variation in high-amplitude network-level co-fluctuations.

### Quotidian variation in edge-level co-fluctuations and hormone concentrations are correlated

In the previous section, we demonstrated that day-to-day variation in gonadotropin concentration was linked to the frequency with which particular high-amplitude “states” are expressed. In that analysis, any co-fluctuation pattern assigned a given community label was treated as a recurrence of the same state. In reality, however, the whole-brain patterns classified by these state motifs exhibited variability between days and even within instances during the same scan. Here, we demonstrate that within-state variation in edge co-fluctuation magnitude is linked not only with gonadotropin concentration, but also with the concentrations of sex hormones estradiol and progesterone.

To link edge-level co-fluctuations with hormones, we needed estimates of communities 1 and 2 for each scan session. To do this, we identified all co-fluctuation patterns assigned to a given community on each scan session and averaged those. Note that in the case of community 1, there was one scan in which it never appeared; in the case of community 2, there were two. We omitted these scans from all analyses carried out in this section. For each of the remaining scans, we created a representative version of community 1 and 2 centroids by averaging all co-fluctuation assigned to that community. We also kept track of the number of samples used to compute the representative pattern (i.e. the number of times that a given state was present in each scan) and after aggregating across all scans, regressed out this number from each node pair, retaining the residuals and calculating their Spearman rank correlation with the concentrations of progesterone, estradiol, luteinizing hormone, and follicle stimulating hormone.

Mass-univariate edge-level analyses can lack statistical power to resolve certain effects. Here, for instance, we performed 79,800 tests (the number of edges) with the aim of identifying those that pass a criterion for statistical significance (for visualization only, we show the edges with the strongest positive and negative correlations embedded in anatomical space in Fig. 4a-d). An alternative strategy is to perform statistical testing at the level of brain systems or communities. Because nodes are aggregated by community, this approach necessarily limits one’s ability to resolve focal, regional effects. However, because the number of comparisons is reduced by an order of magnitude or more, statistical power increases.

**FIG. 4.**
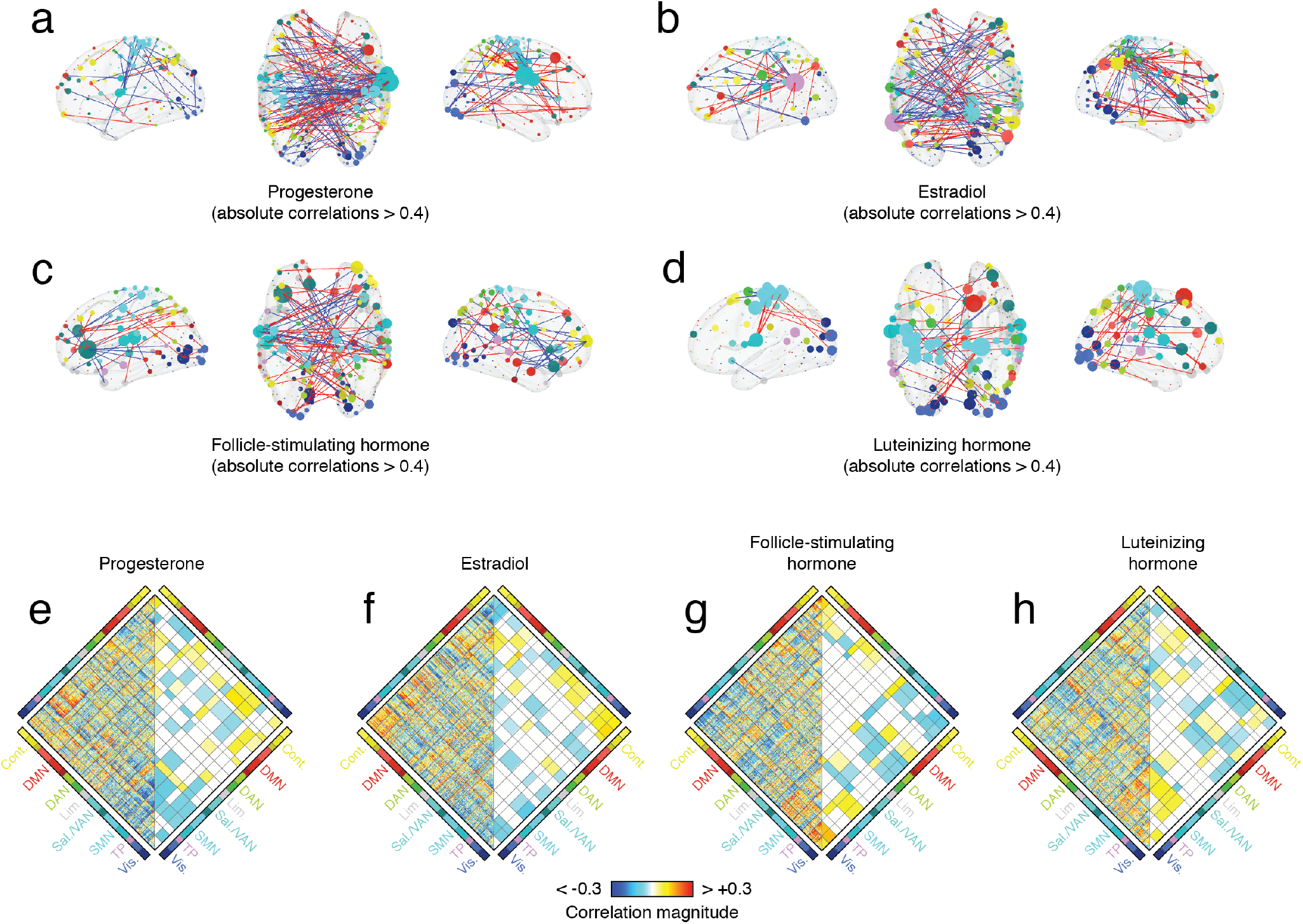
Edge- and system-level correlations with hormone concentration. We calculated the correlation of hormone concentrations with edge-level co-fluctuation magnitudes for cluster 1. Panels *a*-*d* depict strongest correlations in anatomical space. Node size is proportional to the mean correlation magnitude of a node’s edges. Node color was determined by brain system. Edge color denotes positive (red) and blue (negative) correlations. Panels *e*-*h* depict full matrices of correlations (*left*) alongside mean system-level correlations (*right*).

Here, we aggregated edge-level correlations by brain system [31], transforming a correlation matrix of dimensions *ρ*_*region*_ ∈ [400 × 400] into a *ρ*_*system*_ ∈ [16 × 16] matrix. The elements of *ρ*_*system*_ represented the mean correlation coefficients between all pairs of regions assigned to any two systems. We repeated this procedure for all four hormones (Fig. 4e-f), yielding four system-level correlation maps. To identify significant correlations, we repeated the aggregation procedure after randomly permuting system labels using a “spin” test to approximately preserve spatial dependencies between regions (1000 permutations) [32].

In general, we found that the regional correlation patterns for sex hormones were similar to one another (*r*_*P,E*_ = 0.27; *p* < 0.05). The same was true for gonadotropins (*r*_*FSH,LH*_ = 0.38; *p* < 0.05) (Fig. 5a). Due to their similarity and for ease of description, we combined system-level correlation patterns for progesterone with estradiol and FSH with LH, focusing on shared effects (these combined patterns are anti-correlated with one another; *p* < 0.05; Fig. 5b). To summarize shared effects, we classified every pair of systems based on the concordance of correlation patterns. That is, whether both, one, or neither hormones exhibited effects in the same direction (false discovery rate fixed at *q* = 0.05 resulting in adjusted critical value of *p*_*adj*_ = 0.0036; Fig. 5c,d).

**FIG. 5.**
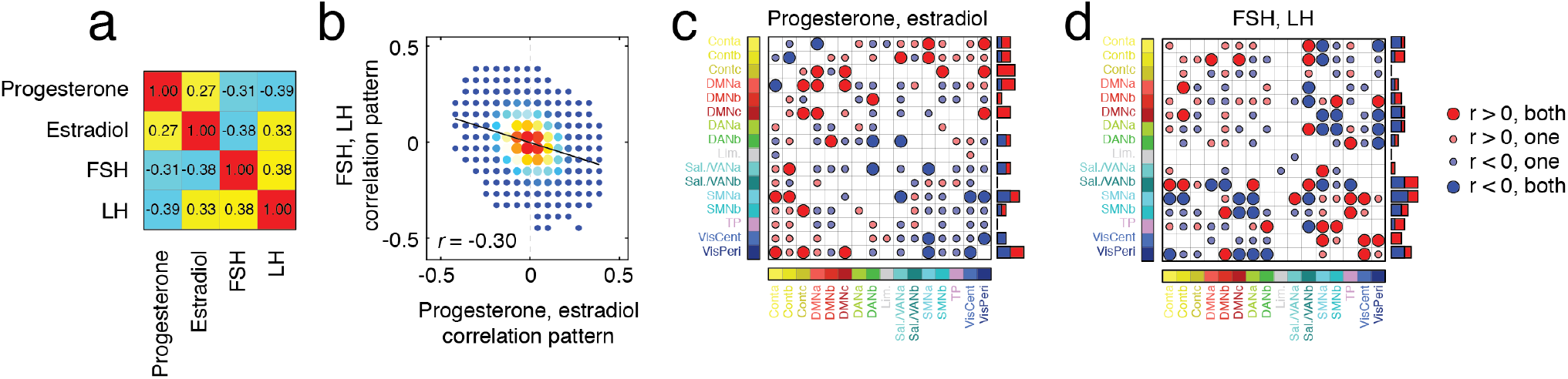
Edge- and system-level correlations with hormone concentration. (*a*) Similarity of correlation patterns between different hormones. (*b*) Two-dimensional histogram showing similarity of correlation patterns from combined sex hormones (progesterone + estradiol) and gonadotropins (FSH + LH). (*c*) Concordance of significant correlations between progesterone and estradiol system-level correlations. Large and small circles indicate high and low levels of concordance. Red and blue colors indicate positive and negative correlations, respectively. The barplot to the right of the matrix is a count of the total number of high concordance interactions in which a given system interacts. Panel *d* depicts analogous information for combined gonadotropins (FSH + LH).

In both cases, we found evidence of broad, brain-wide constellations formed by significant system-level correlations. In the case of sex steroid hormones, correlations involving the control and default mode networks tended to be positive, while correlations involving attention, somatomotor, and visual network tended to be negative (Fig. 5c). On the other hand, in the case of gonadotropins, positive and negative correlations were more uniformly distributed across brain systems, with salience/ventral attention, somatomotor, and visual networks ranking among the systems with the greatest overall number of significant correlations (Fig. 5d).

Collectively, these findings suggest that the co-fluctuation between specific pairs of brain regions are associated with concentrations of sex steroid hormones as well as gonadotropins. These relationships are expressed through distributed, brain-wide constellations of edges linking together many brain systems.

## DISCUSSION

Here, we built upon previous analyses of a dense-sampling dataset in which a single participant underwent daily MRI scans and serological sampling over the course of two full menstrual cycles [23–26]. Leveraging “edge time series” [14, 15], we detected high-amplitude events in each scan session, clustering them into two large communities. The first community reflects opposed activation of default mode and control regions with attentional and sensorimotor regions. We found that the frequency with which it appears across scan sessions was strongly coupled to quotidian fluctuations in the gonadotropic hormones, follicle-stimulating hormone and luteinizing hormone. More generally, we find that variability in the precise co-fluctuation of events assigned to the first community was linked to all four hormones at the level of connections. Our work sets the stage for future studies to investigate relationships between fast fluctuations in network organization and hormones.

### High-amplitude co-fluctuations are linked to endocrine system

Previous applications of edge time series analysis to functional imaging data have reported brief, intermittent, and high-amplitude “events” [14, 16–18, 33]. These studies have shown that the co-fluctuation patterns expressed during events contribute disproportionately to the time-averaged pattern of FC, are subject-specific, can be clustered into a small number of putative “states”, and strengthen brain-behavior associations. For instance, in [14], the authors show that the overall magnitude of brain-behavior correlations increases when the brain measures are derived for high-amplitude co-fluctuations compared to low-amplitude. However, the physiological underpinnings of high-amplitude co-fluctuations remain unclear.

Here, we present evidence that both the frequency with which high-amplitude states occur as well as their topology are strongly correlated with day-to-day fluctuations in hormone concentration. These results suggest that the endocrine system, and specifically reproductive hormones, are a key factor in modulating “events”. While this observation is in line with previous studies that have reported links between reproductive hormones and both brain activity [21, 22] and functional connectivity [23, 26, 34–40], our results extend this link to ultrafast network dynamics. Changes in network structure at this timescale have been relatively unexplored in previous studies due to their cross-sectional study design and low temporal resolution of sliding-window methods for estimating time-varying functional connectivity [41] (although, see [24]).

While the design of our study is not well-suited to support claims about causality, we speculate that by modulating receptor density during the menstrual cycle, hormones may shift the sensitivity of neural circuits, yielding brain states that are increasingly excitable and more likely to produce high-amplitude events [42–44]. Indeed, recent studies have demonstrated that the sex hormones estrogen and progesterone differentially influence the density of steroid hormone receptors across the brain throughout the menstrual cycle [45]. Moreover, this view aligns with the nonlinear and nonstationary contribution of luteinizing hormone to hormone dynamics across the menstrual cycle; large surges in LH near ovulation disrupt the phasic coupling of sex steroid and gonadotropic hormones [41], suggesting that structural changes caused by acute changes in hormone concentration, such as receptor density, influence network connectivity in addition to the direct effects of hormones on cells. Sex hormones are intrinsically involved in micro-level structural changes as well, given that estrogen- and progesterone-mediated changes in subcortical receptor density induce the release of gonadotropin-releasing hormone to induce luteinizing hormone spike [46].

Another possible explanation for why fluctuations in reproductive hormones are associated brain network dynamics concerns their role as neurotransmitters. Previous studies have established links between neurotransmitters and connectivity, demonstrating that regional variation in the concentration of neurotransmitter-related neurons modulate large-scale brain activity, shaping patterns of connectivity and the spontaneous emergence of resting-state networks [47]. For example, dopamine and serotonin oppositionally influence anticorrelations between large-scale networks, such that dopamine is associated with increases and decreases in functional connectivity of the somatomotor and default mode, respectively, while serotonin is associated with the opposite [48]. Other studies have found that neurotransmitter effects on functional connectivity are connected to regional differences in excitatory-inhibitory receptor and hormone ratios [49]. Reproductive hormones, although not traditionally viewed as neurotransmitters, can nonetheless pass through the blood-brain barrier and are known to impact brain function, including memory and anxiety-level behavior [50]. These observations, combined with the fact that edge time series are mathematically exact decompositions of functional connectivity into time-varying contributions, opens the possibility for reproductive hormone to impact patterns of connectivity across time.

We note, however, that while our findings suggest a hormonal contribution to high-amplitude events, other factors likely also play important roles, including the underlying anatomical connectivity, whose network organization shapes the spatial topography of event cofluctuations [17]. Future work should be directed to tease apart the contributions of structural connectivity [51] as well as other factors, including those molecular, vascular, [52] and hemodynamic [53].

### Relationship with results from previous analyses of same data

Here, we make several observations that relate to results from previous studies. First, we found that the state most strongly associated with day-to-day changes in hormone concentrations was typified by opposed cofluctuations of regions in the default mode and control networks with regions in sensorimotor and attentional networks. The coupling of default mode cofluctuations with hormone variation is consistent with results from previous analyses of these same data showing that estradiol fluctuations predict dynamic shifts in default mode network integration in a time-lagged fashion, and that its community structure reconfigures concurrently with spikes in estradiol, luteinizing hormone, and follicle-stimulating hormone across the ovulation window [24, 26].

Second, we find that patterns of edge- and system-level correlations are dissimilar between sex and gonadotropic hormones, but similar within category. While these results suggest differential contributions of sex and gonadotropic hormones to high-amplitude co-fluctuation patterns, they also seemingly contradict results from previous studies [26], which reported opposed effects of estradiol and progesterone.

We note, however, there are some important differences between our study and those previous studies. In particular, our study focuses on rapid, i.e. framewise, changes in co-fluctuation patterns. Even in sliding-window analyses, access to this timescale is limited and the networks resulting from each technique exhibit dissimilar topological profiles [33]. In addition, our study focused on gonadotropic *and* ovarian hormones, and focused not on distinct phases of the menstrual cycle or on peaks in serum levels, but rather on day-to-day fluctuations in hormone levels.

### Future directions

The results of our study present several opportunities for future investigations. First, while our study presents a link between high-amplitude co-fluctuations and female reproductive hormones, its implications for cognitive and clinical processes have not been fully explored. For instance, the original studies also collected behavioral data on the participant’s level of stress, sleep, and affect across the menstrual cycle. Future studies should investigate the role of brain-hormone coupling in mediating behavioral effects.

To our knowledge, this study is the first to our knowledge to identify a significant relationship between luteinizing hormone and daily changes in high-amplitude co-fluctuations in brain connectivity across the menstrual cycle. Many studies have demonstrated luteinizing hormone’s negative association with cognition, and increased luteinizing hormone in females after menopause is associated with poorer cognition and a higher risk of dementia [54, 55]. Future studies should examine how luteinizing hormone’s modulatory effects on cognition relate to daily changes in brain network organization and connectivity in both pre-menopausal and post-menopausal females.

While the dense-sampling framework allows for detailed analyses of single individuals or small cohorts of individuals [27–29], its design makes generalizing to larger and more variable populations challenging. Future studies should analyze relationships between network organization, hormone concentrations, and cognition within subjects across many days of their cycle and between subjects in different endocrine states. Variability in sex hormone production occurs across the female lifespan, starting at the onset of puberty and continuing across the reproductive cycle, pregnancy, and the menopausal transition. Targeting these major neuroendocrine transition states could yield greater insight into sex steroid hormones’ influence on the brain’s network architecture, mood, and cognition. Throughout the life course, changes in women’s reproductive status (e.g. puberty, use of hormonal contraceptives, pregnancy/postpartum, and perimenopause) have been associated with increased risk for mood disturbance including major depressive disorder. Yet, the neurobiological pathways by which endocrine changes give rise to depressive symptoms in some women, but not others, is unclear. A more detailed understanding of how sex hormone fluctuations produce rapid changes in brain network structure could provide a framework for understanding risk and resilience to mood disorders.

### Limitations

Although these two studies provide ample information about daily changes in brain activity and hormone levels, the single-subject nature of the study limits the generalizability of our results. It is possible that the relationships between community organization and hormone concentrations observed in this study are a result of individual processes in the subject that are not present in most of the population. Further deep-phenotyping studies with larger cohorts should be conducted to determine the generalizability of the results across the general population.

A second limitation concerns the detection and characterization of edge time series and high-amplitude cofluctuations. Recent studies have suggested that some of their properties can be anticipated from the static, i.e. time-invariant, functional connectivity matrix alone [56], downplaying their interpretation as dynamic events. Determining the features of higher-order network constructs like edge time series and edge connectivity remains an active area of research [18, 19, 33]. Additional studies are necessary to address this and related open questions.

A final limitation concerns our interpretation of endocrine effects on recorded brain activity and connectivity. Here, we find that hormone concentrations are associated with high-amplitude co-fluctuations that have been shown to shape whole-brain patterns of functional connectivity [14, 16–18, 33]. However, hormones also impact brain vasculature [57]. Because the fMRI BOLD signal is an indirect measure of brain activity that depends critically on neurovascular coupling, an alternative explanation is that true effect of gonadotropins is on brain vasculature, which modulates the hemodynamic response. Future, and more targeted, studies should aim to disentangle these effects.

### Conclusions

In conclusion, our study posits a link between high-amplitude, network-level co-fluctuations and the human endocrine system. Specifically, we report an association between the frequency of dynamic network states and variation in luteinizing and follicle-stimulating hormones. Our work addresses questions concerning the factors contributing to high-amplitude co-fluctuations while opening up new opportunities for future studies

## MATERIALS AND METHODS

### Datasets

Neuroimaging and endocrine data comes from a single subject (author L.P.) scanned over a course of 30 days, on two separate occasions (Study 1 and Study 2). The subject had no history of neuropsychiatric diagnosis, endocrine disorders, or prior head trauma and no history of smoking. She had a history of regular menstrual cycles (no missed periods, cycle occurring every 26–28 days). In the 12 months prior to the first 30-day data collection period, the subject had not taken hormone-based medication. In the second study, the participant was on a hormone regimen (0.02mg ethinyl-estradiol, 0.1mg levonorgestrel, Aubra, Afaxys Pharmaceuticals), which she began 10 months prior to the start of data collection. The pharmacological regimen used in Study 2 chronically and selectively suppressed progesterone while leaving estradiol dynamics largely indistinguishable from Study 1. The participant gave written informed consent and the study was approved by the University of California, Santa Barbara Human Subjects Committee.

Neuroimaging data was collected on a Siemens 3T Prisma scanner with a 64-channel phased-array head coil. Scans were collected around the same time each day (11:00 am local time). For each scanning session, a highresolution T1-weighted anatomical sequence (MPRAGE) was acquired (TR = 2500 ms, TE = 2.31 ms, T_1_ = 934 ms, flip angle = 7°). Following this, a 10-minute resting state sequence was acquired, using a T_2_*-weighted multiband echo-planar (EPI) sequence was acquired (72 oblique slices, TR = 720 ms, TE = 37 ms, voxel size 2 mm^3^, flip angle = 52°, multiband factor = 8). To mitigate against motion, a custom 3D-printed foam head case was employed for days 8-30 of the first 30-day period, and for days 1-30 of the second 30-day period.

### Endocrine procedures

A licensed phlebotomist inserted a saline-lock intravenous line into the dominant or non-dominant hand or forearm daily to evaluate hypothalamic-pituitary-gonadal axis hormones, including serum levels of gonadal hormones (17*β*-estradiol, progesterone, and testosterone) and the pituitary gonadotropins luteinizing hormone (LH) and follicle stimulating hormone (FSH). One 10 cc mL blood sample was collected in vacutainer SST (BD Diagnostic Systems) each session. The sample clotted at room temperature for 45 minutes until centrifugation (2000×g for 10 minutes) and were then aliquoted into three 1 mL microtubes. Serum samples were stored at −20 ° until assayed. Serum concentrations were determined via liquid chromatography mass-spectrometry (for all steroid hormones) and immunoassay (for all gonadotropins) at the Brigham and Women’s Hospital Research Assay Core. Assay sensitivities, dynamic range, and intra-assay coefficients of variation (respectively) were as follows: estradiol 1 pg/mL, 1-500 pg/mL, < 5% relative standard deviation (RSD), 0.05 ng/mL, 0.05-10 ng/mL, 9.33% RSD; testosterone, 1.0 ng/dL 1-2000 ng/dL, < 4% RSD. FSH and LH levels were determined via chemiluminescent assay (Beckman Coulter). The assay sensitivity, dynamic range, and intra-assay coefficient of variation were as follows: FSH, 0.2 mlU/mL, 0.2-200 mIU/mL, 3.1-4.3%; LH, 0.2 mIU/mL, 0.2-250 mIU/mL, 4.3-6.4%. Importantly, we note that LC-MS assessments of exogenous hormone concentrations (attributable to the hormone regime itself) showed that serum concentrations of ethinyl estradiol were very low (*M* =0.01 ng/mL; range 0.001-0.016 ng/mL) and below 1.5 nl?mL for levonorgestrel (*M* = 0.91 ng/mL; range = 0.03-1.45 ng/mL). This ensures that the brain hormone associations reported in Study 2 are still do to endogenous estradiol action.

#### Image preprocessing

Preprocessing was performed using *fMRIPrep* (20.2.0) [58], which is based on Nipype 1.5.1 [59]. The following description of fMRIPrep’s preprocessing is based on boilerplate distributed with the software covered by a ‘no rights reserved’ (CCO) license. Internal operations of fMRIPrep use Nilearn 0.6.2 [60], mostly within the functional processing workflow. The preprocessing pipelines employ functions from ANTs (2.3.3), FreeSurfer (6.0.1), FSL (5.0.9), and AFNI (20160207).

T1-weighted (T1w) images were corrected for intensity non-uniformity (INU) with `N4BiasFieldCorrection` [61], distributed with ANTs [62], and used as T1wreference throughout the workflow. The T1w-reference was then skull-stripped with a Nipype implementation of the `antsBrainExtraction.sh` workflow, using NKI as target template. Brain tissue segmentation of cere-brospinal fluid (CSF), white-matter (WM) and graymatter (GM) was performed on the brain-extracted T1w using `fast` [63]. Brain surfaces were reconstructed using `recon-all` [64], and the brain mask estimated previously was refined with a custom variation of the method to reconcile ANTs-derived and FreeSurfer-derived segmentations of the cortical gray-matter of Mindboggle [65]. Volume-based spatial normalization to one standard space (MNI152NLin2009cAsym) was performed through nonlinear registration with `antsRegistration`, using brain-extracted versions of both T1w reference and the T1w template. The following template was selected for spatial normalization: *ICBM 152 Nonlinear Asymmetrical template version 2009c* [66]. For each of the BOLD runs found per subject (across all tasks and sessions), the following preprocessing was performed. First, a reference volume and its skull-stripped version were generated using a custom methodology of *fMRIPrep*. Susceptibility distortion correction (SDC) was omitted. The BOLD reference was then co-registered to the T1w reference using `bbregister` (FreeSurfer) which implements boundary-based registration [67]. Co-registration was configured with six degrees of freedom. Head-motion parameters with respect to the BOLD reference (transformation matrices, and six corresponding rotation and translation parameters) are estimated before any spatiotemporal filtering using `mcflirt` [68]. BOLD runs were slice-time corrected using `3dTshift` from AFNI [69]. The BOLD time-series (including slice-timing correction when applied) were resampled onto their original, native space by applying the transforms to correct for head-motion. These resampled BOLD time-series will be referred to as *preprocessed BOLD in original space*, or just *preprocessed BOLD*. The BOLD time-series were resampled into standard space, generating a *preprocessed BOLD run in MNI152NLin2009cAsym space*. First, a reference volume and its skull-stripped version were generated using a custom methodology of fMRIPrep. Several confounding time-series were calculated based on the preprocessed BOLD: framewise displacement (FD), DVARS and three region-wise global signals. FD was computed using two formulations following Power (absolute sum of relative motions, [70] and Jenkinson (relative root mean square displacement between affines, [68]). FD and DVARS are calculated for each functional run, both using their implementations in Nipype following the definitions by[70]. The three global signals are extracted within the CSF, the WM, and the whole-brain masks. Additionally, a set of physiological regressors were extracted to allow for component-based noise correction [71]. Principal components are estimated after high-pass filtering the *preprocessed BOLD* time-series (using a discrete cosine filter with 128s cut-off) for the two *CompCor* variants: temporal (tCompCor) and anatomical (aCompCor). tCompCor components are then calculated from the top 2% variable voxels within the brain mask. For aCompCor, three probabilistic masks (CSF, WM and combined CSF+WM) are generated in anatomical space. The implementation differs from that of Behzadi et al. in that instead of eroding the masks by 2 pixels on BOLD space, the aCompCor masks are subtracted a mask of pixels that likely contain a volume fraction of GM. This mask is obtained by dilating a GM mask extracted from the FreeSurfer’s *aseg* segmentation, and it ensures components are not extracted from voxels containing a minimal fraction of GM. Finally, these masks are resampled into BOLD space and binarized by thresholding at 0.99 (as in the original implementation). Components are also calculated separately within the WM and CSF masks. For each CompCor decomposition, the *k* components with the largest singular values are retained, such that the retained components’ time series are sufficient to explain 50 percent of variance across the nuisance mask (CSF, WM, combined, or temporal). The remaining components are dropped from consideration. The head-motion estimates calculated in the correction step were also placed within the corresponding confounds file. The confound time series derived from head motion estimates and global signals were expanded with the inclusion of temporal derivatives and quadratic terms for each [72]. Frames that exceeded a threshold of 0.5 mm FD or 1.5 standardised DVARS were annotated as motion outliers. All resamplings can be performed with *a single interpolation step* by composing all the pertinent transformations (i.e. head-motion transform matrices, susceptibility distortion correction when available, and co-registrations to anatomical and output spaces). Gridded (volumetric) resamplings were performed using `antsApplyTransforms`, configured with Lanczos interpolation to minimize the smoothing effects of other kernels [73]. Non-gridded (surface) resamplings were performed using `mri_vol2surf` (FreeSurfer).

#### Parcellation

The Schaefer parcellation [31] was used to delineate 400 regions on the cortical surface. To transfer the parcellation from *fsaverage* to *subject* space, FreeSurfer’s `mris_ca_label` function was used in conjunction with a pre-trained Gaussian classifier surface atlas [74] to register cortical surfaces based on individual curvature and sulcal patterns. The result is a volumetric parcellation rendered in subject anatomical space that follows the cortical ribbon estimated via FreeSurfer’s `recon-all` process.

#### Functional Connectivity

Each preprocessed BOLD image was linearly detrended, band-pass filtered (0.008-0.08 Hz), confound regressed and standardized using Nilearn’s signal.clean function, which removes confounds orthogonally to the temporal filters. The confound regression strategy included six motion estimates, mean signal from a white matter, cerebrospinal fluid, and whole brain mask, derivatives of these previous nine regressors, and squares of these 18 terms [72]. Spike regressors were not applied. The 36 parameter strategy (with and without spike regression) has been show to be a relatively effective option to reduce motion-related artifacts [75]. An alternative preprocessesing strategy was also employed to evaluate the stability of the findings. This strategy included six motion estimates, derivatives of these previous six regressors, and squares of these 12 terms, in addition to five anatomical CompCor components [71]. Following these preprocessing operations, the mean signal was taken at each node in volumetric anatomical space.

### Edge time series

Let *z*_*i*_ = [*z*_*i*_(1), … *z*_*i*_(*T*)] be the z-scored time series for region *i*. Most network neuroscience analyses define the functional connectivity between pairs of regions {*i, j*} as the Pearson correlation of their activity, i.e. 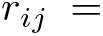 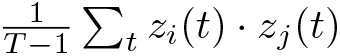.

Recently, we proposed a “temporal unwrapping” of functional connection weights by omitting the summation. That is, we calculate the instantaneous magnitude and sign of co-fluctuation between pairs of brain regions as *e*_*ij*_(*t*) = *z*_*i*_(*t*) · *z*_*j*_(*t*) [14, 15, 18]. This approach has the benefit of resolving changes in pairwise interactions at a temporal resolution of single frames. It also is deeply related to the Pearson correlation and functional connectivity – the temporal average of a region pairs’ edge time series is exactly equal to its connection weight. In this way, edge time series can be viewed as precise temporal decompositions of functional connectivity. In this study, we calculated edge time series for every pair of cortical and subcortical regions for every scan session.

### Event detection

The brainwide level of network activity at any instant can be summarized as the root sum of square over all edges co-fluctuations:

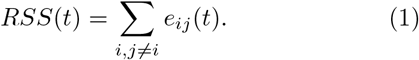

In previous studies we demonstrated that *RSS* exhibits “bursty” behavior, such that most time points express low-amplitude co-fluctuations while a relatively small number exhibit large *RSS* values [18]. These “events” are thought to reflect underlying anatomical connectivity [17], are highly individualized [18], and contribute disproportionately to the time-averaged pattern of functional connectivity [14, 16].

Recently, we proposed a simple statistical test for detecting events. This test works by comparing observed *RSS* values with those generated under a temporal null model in which regional time series are circularly-shifted by a random offset in either direction. This procedure generates a null distribution of *RSS* values against which an empirical value can be compared statistically. For each frame we estimated a non-parametric p-value by counting the fraction of null *RSS* values that exceeded the observed value. We compared p-values against an adjusted critical value while fixing the false discovery rate at *q* = 0.05 [76]. Note that this null model necessarily destroys the correlation structure (functional connectivity) and can be viewed as a principled method for selecting a threshold for events.

Once a statistical threshold for events is determined, we identify temporal contiguous sequences of suprathreshold frames, which we refer to as *event segments*. We discard any such segments that include a high-motion frame or are within two frames of a high-motion frame and, from those segments, extract as a representative cofluctuation pattern the frame corresponding to the peak *RSS*. We repeat this procedure for all scans, retaining the peak co-fluctuation patterns.

### Cluster definition

In previous studies, we demonstrated that high-amplitude co-fluctuations patterns can be clustered into a series of states [18]. To do this, we compute the similarity (correlation) between all pairs co-fluctuation patterns extracted during event segments. This results in a pattern × pattern matrix, which we submitted to a generalized version of the Louvain algorithm [77, 78] for modularity maximization algorithm [79]. Modularity maximization is a computational heuristic for detecting community structure in networked data. It defines communities (clusters) as groups elements whose internal density of connections maximally exceed what would be expected. In this context, we defined the expected weight of connections to be equal to the mean similarity between all pairs of patterns.

Modularity maximization with the Louvain algorithm is non-deterministic and, depending upon initial conditions, can yield dissimilar results. Accordingly, we ran the algorithm 1000 times with different random seeds. We resolved variability across these different seeds using a consensus clustering algorithm in which we iteratively clustered the module co-assignment matrix until convergence (see any of [18, 80, 81] for details of this algorithm). The resulting consensus partition assigned each co-fluctuation pattern to non-overlapping clusters.

## AUTHOR CONTRIBUTIONS

SG and RFB conceived of project and performed analysis. LP, TS, EGJ contributed data. JF processed data. All authors wrote manuscript.

## ACKNOWLEDGMENTS

We thank Haily Merritt and Jacob Tanner for providing feedback on an early version of this manuscript. This material is based upon work supported by the National Science Foundation under Grant No. 2023985 (RFB).

---

**FIG. S1.**
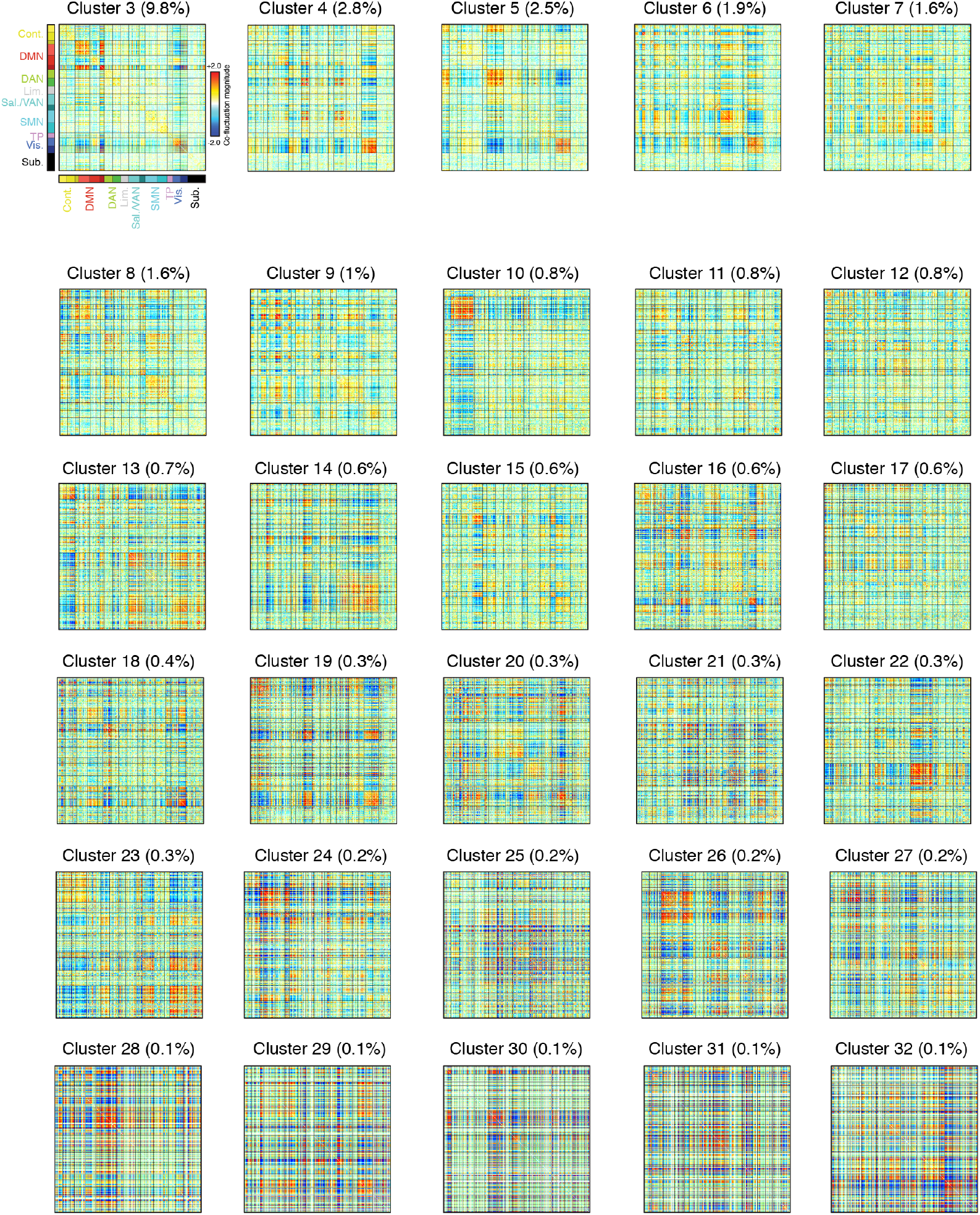
Centroids for remaining co-fluctuation communities. In the main text we clustered “event” co-fluctuation patterns and analyzed the top two communities by frequency. Here, we show the centroids for the remaining 30 communities.

**FIG. S2.**
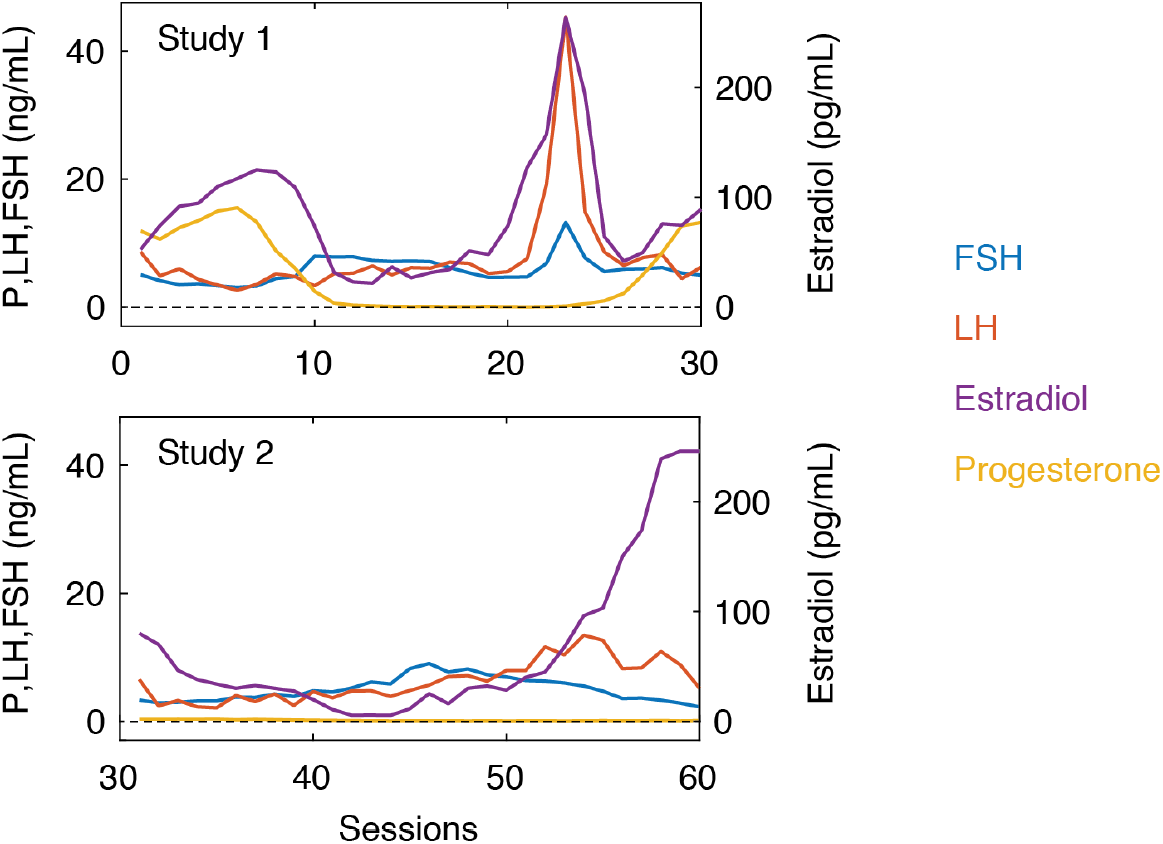
Hormone fluctuations across experimental sessions. Here, we show variation of progesterone, estradiol, follicle-stimulating hormone, and luteinizing hormone across the Study 1 and Study 2 datasets.

**FIG. S3.**
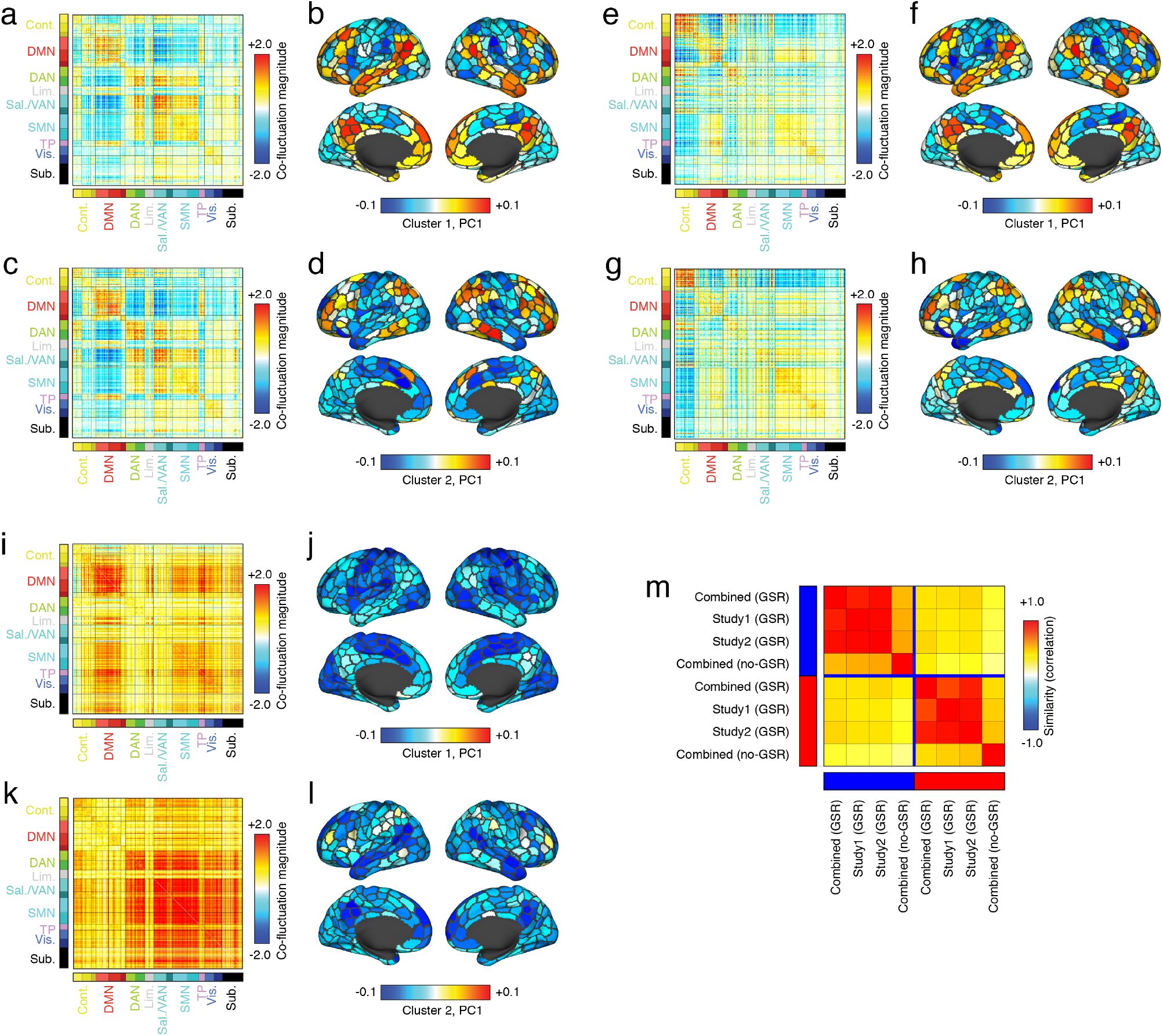
Robustness of communities 1 and 2 to processing decisions. In the main text we clustered “event” cofluctuation patterns and analyzed the top two communities by frequency. To demonstrate the robustness of these two communities, we repeated this procedure, by splitting the 60 scan sessions into their two respective experiments (Study 1 and Study 2; panels *a*-*d* and *e*-*h*, respectively). We also analyzed data processed without global signal regression (panels *i*-*l*). In each quartet of panels, we highlight the top two communities by frequency. In panel *m*, we show the similarity of community centroids to one another. In general, we find that when splitting data by experiment, we maintain an excellent correspondence with the original data. We also find a strong correspondence between the original data and data processed without global signal regression.

**FIG. S4.**
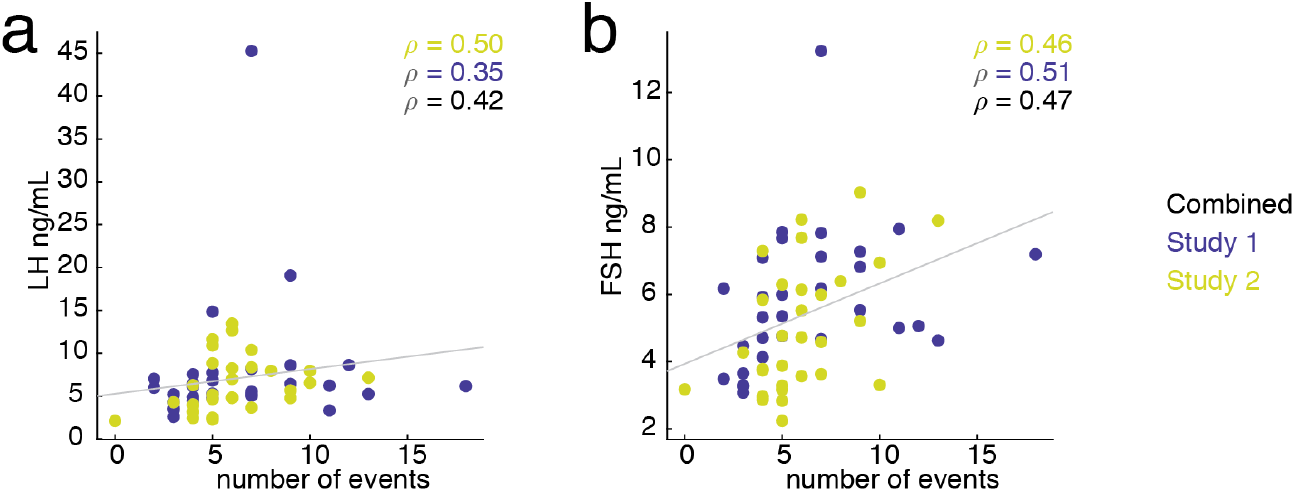
Robustness of correlation between state frequency and gonadotropins. In the main text we reported correlations between gonadotropin concentration and the frequency (counts) of community (brain state) 1 with luteinizing and follicle-stimulating hormone. Here, we repeat the analysis after splitting the data by experiment (Study 1 *versus* Study 2). (*a*) Scatterplot of community frequency and luteinizing hormone (black is combined correlation, yellow is Study 1, blue is Study 2). (*b*) Analogous plot, but for follicle-stimulating hormone.

**FIG. S5.**
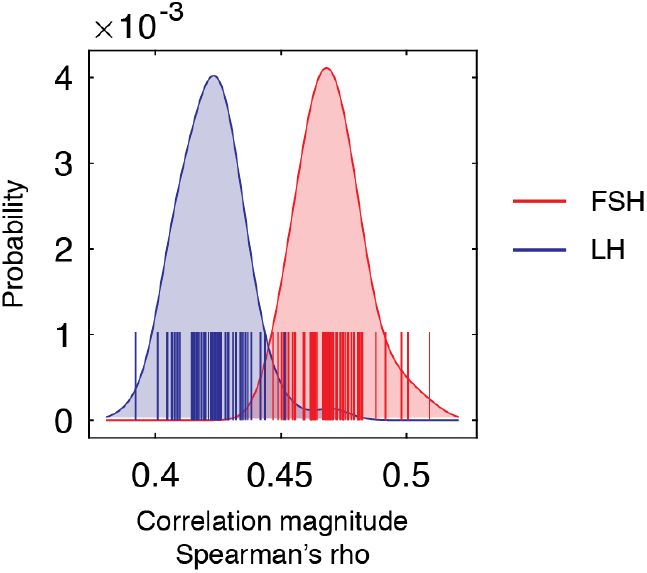
Impact of “leave one out” analysis on state frequency and gonadotropin correlations. Distribution of correlation coefficients after removing data from single scans and recomputing the Spearman correlation coefficient. In this plot, red corresponds follicle-stimulating hormone and blue corresponds to luteinizing hormone.

**FIG. S6.**
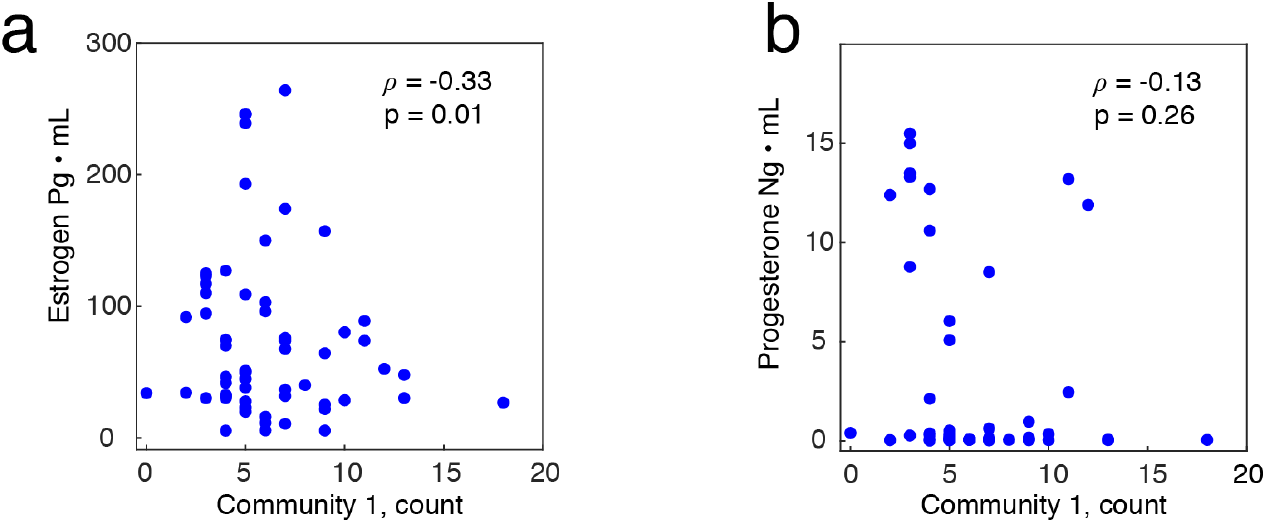
Correlations of sex hormone concentration with frequency of community 1. In the main text, we reported positive correlations between gonadotropic hormones and the frequency of community 1. Here, we report analogous results for sex hormones (*a*) estradiol and (*b*) progesterone. Note that the correlation with estradiol is statistically significant only without correcting for multiple comparisons.

**FIG. S7.**
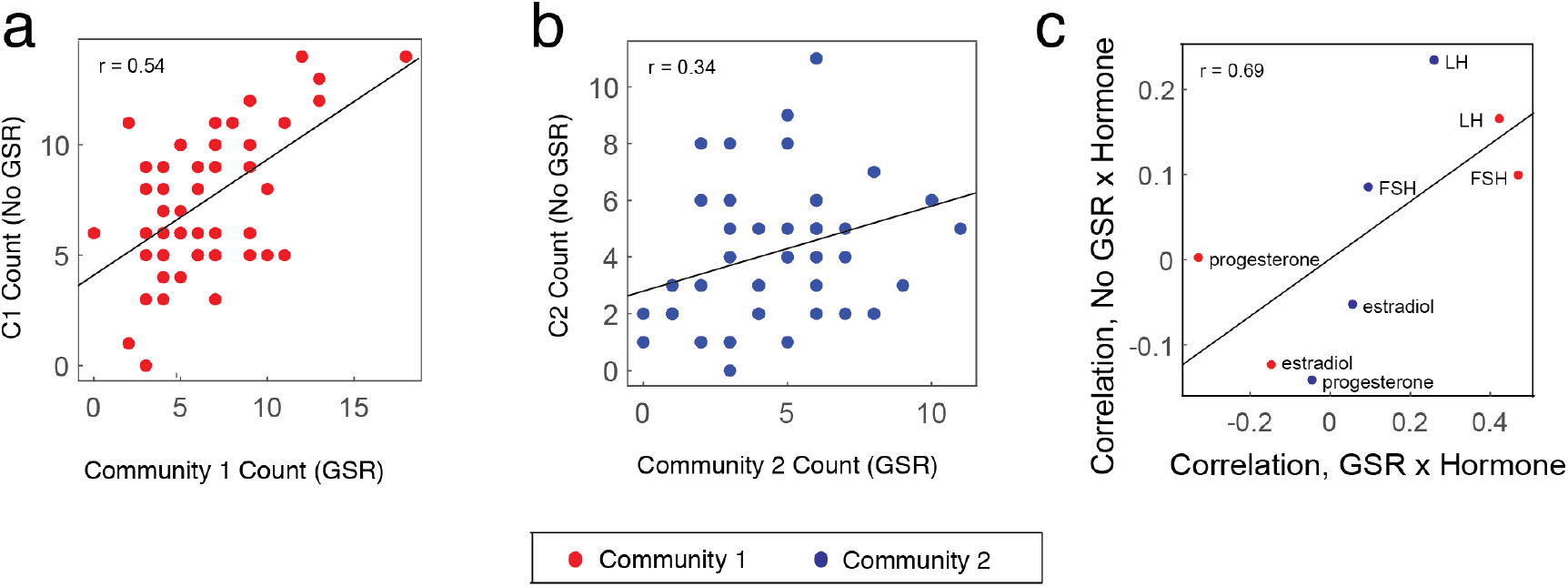
Effect of global signal regression on state frequency and correlations. In the main text we analyzed fMRI data that had been processed using a pipeline that included global signal regression. Here, we report results using the same data but processed without global signal regression. The analysis procedures were performed identically for both datasets. After obtaining consensus clusters, we mapped communities to those obtained following the global signal regression pipeline. Panels *a* and *b* show correlations between state frequencies (how often a given community appeared on any one scan session), for communities 1 and 2. We find a positive correspondence in both cases. In the main text we also computed the correlation between state frequency and hormone concentration. We find that without global signal regression, the overall magnitude of correlations is decrease. However, the overall pattern of correlations is largely preserved (see panel *c*).

## Notes

### Competing Interest Statement

The authors have declared no competing interest.

